# How to make a kin selection model when marginal fitness is non-linear?

**DOI:** 10.1101/007674

**Authors:** R.H. Schonmann, R. Boyd, R. Vicente

## Abstract

We observe that the Taylor-Frank method for making kin selection models when fitness *w* is a nonlinear function of a continuous actor’s phenotype *y* and the average phenotype *z* in its social environment requires *w*(*y, z*) to be differentiable (as a function of two variables, i.e., jointly in *y* and *z*). This means that even if *w*(*y, z*) is non-linear globally, locally it must be close to linear, meaning that its graph must be well approximated by a plane. When more than two individuals interact, this assumption is only satisfied when the marginal fitness of the actor is a linear function of the fraction of individuals in its social environment that share its phenotype. This assumption sometimes fails for biologically important fitness functions, for instance in microbial data and the theory of repeated n-person games. In these cases, the Taylor-Frank methodology cannot be used, and a more general form of direct fitness must replace it, to decide when a social mutant allele can invade a monomorphic population.

## Introduction

According to Hamilton’s rule, the fitness of an allele should be measured by how much it affects the reproductive success of its carriers, added to the effect that its carriers have on the reproductive success of others weighted by relatedness [1]. In its original formulation, Hamilton’s rule required additive fitness effects [2], and a number of extensions have been developed to deal with nonlinearities (reviewed, e.g., in [3–5]). In one of the most influential extensions, [6], it is assumed that (1) fitnesses are functions of two variables: a continuously varying actor’s phenotype *y* and the average value *z* of this phenotype in the actor’s social environment, and (2) phenotypic variation is small. This approach has been applied to a wide range of biological problems (see, e.g., [3], [7], [4, Box 6.1] and [5, Box 6]), and it has been suggested (see e.g., [3, pp. 37-38], [4, pp. 137-138] and [5, Box 6]) that it shows that Hamilton’s rule, in terms of marginal costs and benefits, can always be applied to problems with continuously varying phenotypes, as long as variation in the traits is small enough. However, as we explain in the next section, the Taylor-Frank method depends on the assumption that fitness functions are differentiable as functions of *y* and *z*. (To be clear: differentiable as function of two variables, not just differentiable in each variable separately [8]). This means intuitively that in the relevant region of small phenotypic variation, the fitness function must be close to a linear function of *y* and *z*. When more than two individuals interact, this assumption is only satisfied when the marginal fitness of the actor is a linear function of the fraction of individuals in its social environment that share its phenotype. And in the section on biological significance we point out that fitness functions that occur in real biological applications may not be differentiable (see, e.g., equation (6) that corresponds to an *n*-player repeated interaction). When this is the case one cannot use the chain rule of multi-variable calculus that is the basis of the Taylor-Frank method. Some important treatments of social evolution that utilize the Taylor-Frank method assume explicitly that fitness functions are differentiable as functions of the relevant variables (see, e.g., [9, p. 95]), and in this way restrict the applicability of their conclusions. But others, ([3, pp. 37-38], [4, pp. 137-138] and [5, Box 6] omit the assumption and come to conclusions that are not as widely applicable as they suggest. Moreover, in at least one published paper [10] the Taylor-Frank method was applied in a situation in which the differentiability condition is violated, yielding incorrect expressions for the evolutionary stability of equilibria (see the end of the section on biological significance). We will explain also in the next section, how the Taylor-Frank direct fitness method can be generalized to apply to problems for which one cannot make this assumption of differentiability.

## Invasion by a rare mutant under weak selection

The Taylor-Frank direct fitness approach assumes that the fitness of an individual is affected by its own phenotype and the phenotypes of other individuals in its social environment. Individual phenotype is represented by a heritable quantitative character *y*. For instance, *y* could represent the amount of some costly to produce substance that the individual secretes in the environment and that is beneficial to nearby individuals. Initially all individuals in the population have the same value of this character, 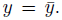 Rare mutations produce a variant with 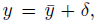 where *|δ|* is small. The rare mutant will invade if its average fitness, *w*m, is higher than that of the wild type, *w*w. To compute these average fitnesses, randomly select a focal individual from the population. Following [6], we denote by *w*(*y, z*) the fitness of a focal individual with phenotype *y*, in a social environment in which the average phenotype is *z* (note that *w*(*y, z*) depends also on 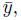 but since this quantity is fixed, it is omited from this notation). If the focal is a wild type, all the individuals in its social environment are likely to be wild types, and therefore, 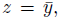 and 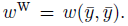 But if the focal individual is a mutant, other individuals in its social environment may also be mutants, for instance due to common descent. Let *X* be the random variable that represents the fraction of members of the social environment of a mutant individual that are mutants. Then 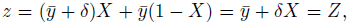 and 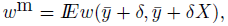 where the expectation (denoted by *IE*) is taken over social environments, i.e., over *X*. The mutants will invade the population when 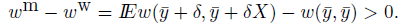 Because *|δ|* is small, one needs only to consider the behavior of the function *w*(*y, z*) in the neighborhood of the point where 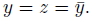 Key to the Taylor-Frank approach is the assumption that the chain rule of multi-variable calculus applies and gives, neglecting terms that are much smaller than *|δ|*,

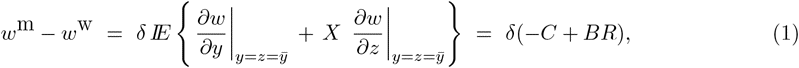

where −*C* and *B* are the values of the partial derivatives in the *y* and *z* directions at the point 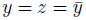 and

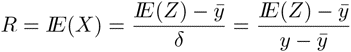

is the average relatedness in the social environment of a mutant individual (see [2]). Provided that one can apply the chain rule, as above, the conclusion is that Hamilton’s rule

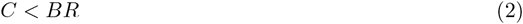

is the necessary and sufficient condition for the mutants to invade the monomorphic population with phenotype 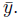

However, the use of the chain rule requires (see, e.g., [8]) that the function *w*(*y, z*) be differentiable, meaning that it is well approximated by a linear function of *y* and *z*, in the neighborhood of 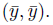 That is, up to an error term that is much smaller than 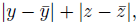 we must have in good approximation *w*(*y, z*) = *α* + *βy* + *γz*, in the neighborhood of 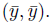 In other words, the surface that represents the function *w*(*y, z*) must be well approximated by a plane in the neighborhood of 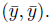 In a one-dimensional setting, differentiability at a given point means that the graph of the function is well approximated by a straight line close to this point. In two dimensions, differentiability at a point means that the graph of the function is well approximated by a plane close to this point. To better understand this well known concept, see its biological meaning and relate it to the chain rule, assume only that for every value of *x* in the range from 0 to 1, the “directional derivative” of *w*(*y, z*),

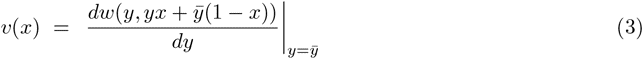

in the direction of the straight line 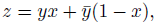 is well defined. (The proper directional derivative is defined as 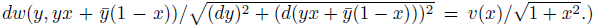 This “directional derivative” *v*(*x*) gives the incremental fitness effect of changes in *y* for a given fixed fraction *x* of mutants in the social environment. These derivatives can exist for every value of *x* and at the same time *w*(*y, z*) can fail to be differentiable in (*y, z*)—see Fig. 1 for an example in which *w*(*y, z*) is not differentiable, despite *v*(*x*) being a smooth function. Neglecting an error term much smaller than *δ*, we have

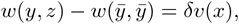

when 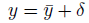 is close to 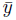 and 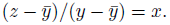 The quantity *v*(*x*) is therefore the marginal fitness of a focal mutant, in a social environment with a fraction *x* of mutants. Differentiability of the function of two variables *w*(*y, z*) at 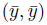 implies, through the chain rule applied to (3), that *v*(*x*) must be the linear function of *x* given by *v*(*x*) = −*C* + *Bx*, with 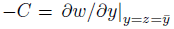 and 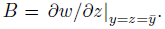 This condition may or not hold in biologically relevant situations (see section on biological significance). It is only when it holds that (1) is valid and (2) is the correct condition for invasion.

This means that the Taylor-Frank approach does not apply to situations in which the marginal fitness *v*(*x*) of the mutants is a non-linear function of the *fraction x* of mutant individuals in the social environment. There is nevertheless no difficulty in obtaining a valid (direct fitness, kin selection) condition for invasion, based only on the assumption that the “directional derivatives” *v*(*x*) exist. The quantity *δv*(*x*) is close to the difference in fitness between a mutant focal individual in a social environment with a fraction *x* of mutant individuals and the fitness of a wild focal individual in an environment in which everyone is of wild type. Therefore, when *δ* is small, neglecting terms that are much smaller than *|δ|*, we have

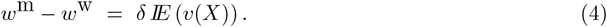

**Figure 1.**
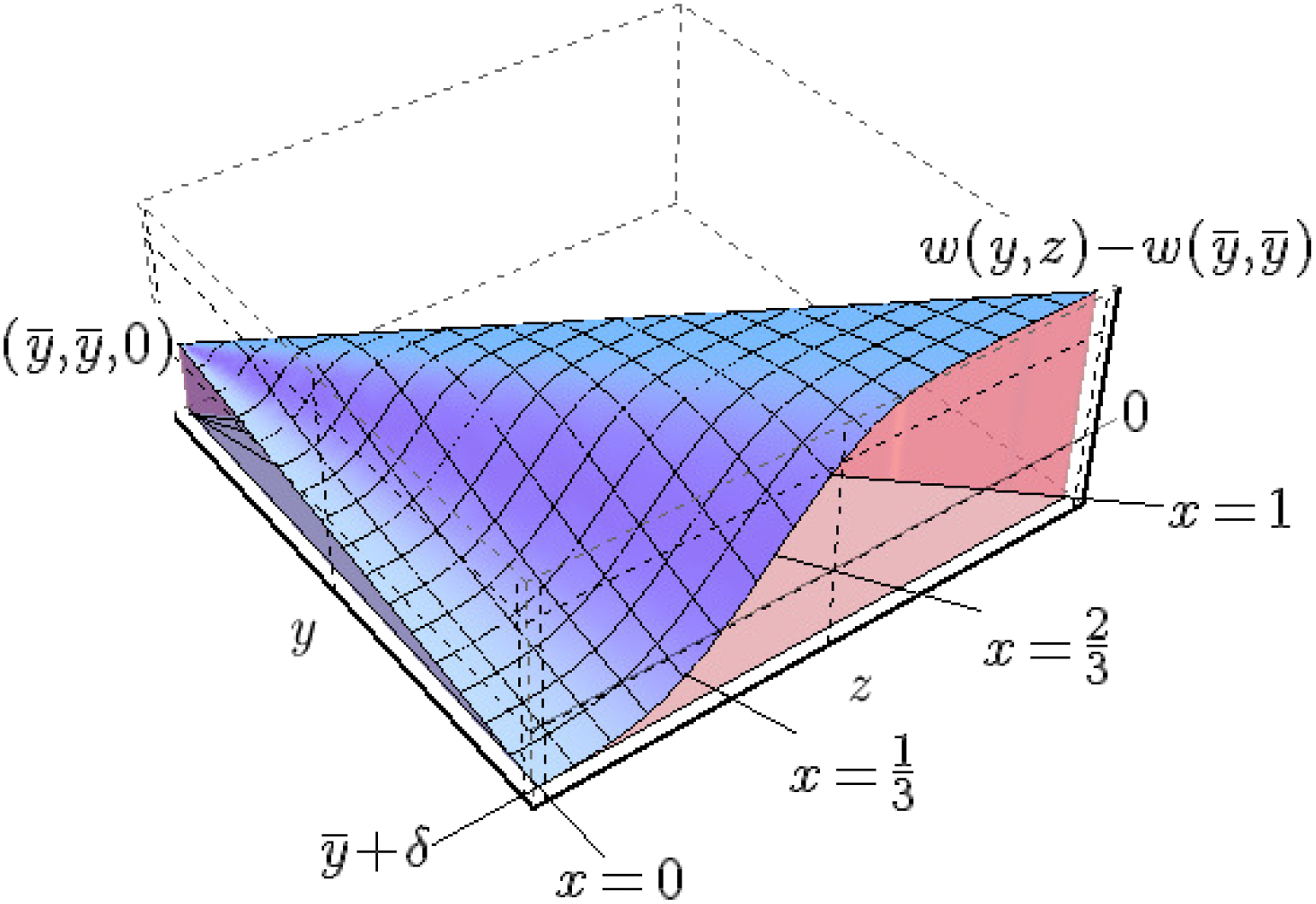
The surface representing 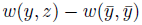 in the neighborhood of the point 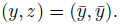 This point appears in the left side of the picture, and the function takes the value 0 there. In the picture *y* ranges from 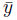 to 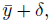 and *z* ranges from 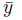 to *y*. The parameter 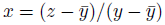 identifies directions in the (*y, z*) plane, away from the point 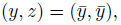 and biologically represents the fraction of individuals in the social environment of a mutant focal individual that are also mutants. The values of the “directional derivatives” *v*(*x*), which represent marginal fitnesses, are indicated by the s-shaped curve produced by the intersection of the surface with the plane 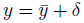 (this s-shaped curve appears as the frontal border of the blue surface in the picture). The surface would only be well approximated by a plane, in the neighborhood of 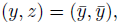 if *v*(*x*) were a linear function, rather than s-shaped. Notice that *w* is not differentiable anytime that *v*(*x*) is a non-linear function of the fraction of mutant-types in the social environment; no kinks or discontinuities are necessary. When, as in this picture, differentiability at 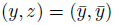 fails, one can not use the chain rule as in the derivation of (1), but the more general (4) still applies and provides the direction of selection.

The condition for the mutants to invade the monomorphic population with phenotype 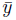 is therefore

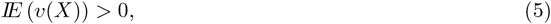

which generalizes Hamilton’s rule (2), and reduces to it precisely when *v*(*x*) = −*C* + *Bx* is a linear function of *x*, i.e., when the interaction of the individuals in each environment affects fitnesses of mutants as a linear public goods game. We will show in a later section that the same conclusion holds when costs and benefits are conceptualized as regression coefficients of fitness against phenotypic value. To compute the expected value in (5) one needs to know the distribution of *X*. In the next section we will discuss cases in which this has been done, and (5) was used to evaluate theory against empirical population parameters.

The contrast between the simplicity and apparent generality in the derivation of (1) and the limitations explained in the previous paragraph may seem puzzling at first sight. This apparent paradox is solved once one understands that condition (1) relies on the assumption that *w*(*y, z*) is differentiable in the neighborhood of the point where 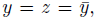 and that this means that the surface that represents this function is well approximated by a plane close to that point. To see why this assumption can fail, consider Fig. 1 in which the difference in the fitness of mutants and wild types, 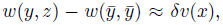 is a sigmoidal function of the fraction of mutants, *x*, for each given value of *δ*. As the value of *δ* decreases, the values of 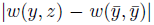 approach 0, but the sigmoidal shape does not change, implying that also the limit *v*(*x*) is sigmoidal, rather than a linear function. This means that the surface representing *w*(*x, z*) cannot be approximated by a plane, close to 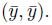 Of course, as the usefulness of the Taylor series approach throughout science attests, most nonlinear functions of interest (in two variables) are well approximated by a plane in the neighborhood of a point. When this happens in our setting, the Taylor-Frank approach is correct. However, as we explain in the next section, there are important biological applications for which data or theory indicate that this is not the case and (1) and (2) are not a good approximation for (4) and (5), which instead are proper expressions of kin selection in those cases (assuming rarity of the mutant type and small trait variability, i.e., small *|δ|*).

It is important to observe that (5) is the appropriate condition for invasion, regardless of the size of the social environment. The random variable *X* only takes values that are multiples of the inverse of the size of the social evironment and that lie between 0 and 1. In the extreme case of diadic interactions, the social environment has a single individual and *X* can only take the values 0 or 1. Since functions with only two points in their domain are always linear, this means that in this case (5) reduces to (2). For social environments with more than 2 individuals *v*(*x*) may or not be linear, and it is precisely when it is linear that (5) reduces to (2), and not otherwise (see [11] for a similar discussion).

## Biological significance

There are at least two biological contexts in which non-linear marginal fitness functions *v*(*x*) are important.

First, experimental evidence from micro-organisms [12–14] is available, for instance from direct manipulation of the fraction of individuals with different genotypes/phenotypes in a social environment and measurements of their rates of reproduction. Such data indicates that sometimes *v*(*x*) is non-linear in the fraction of mutants *x*. One could argue that experimental data does not refer to a limit in which *δ →* 0, and that the data comes from situations in which selection may be strong. This raises the question of how small *δ* has to be for one to regard selection as weak. Basically selection is weak when the differences in phenotype in the population produce only minor differences in reproductive success, so that one can compute (4) assuming that the expectation corresponds to neutral drift without selection. (Separation of time scales; see, e.g., [9, 15–17].) Whether *δ* can be that small while *v*(*x*) is empirically non-linear is an important question to be investigated experimentally.

Second, in repeated n-player games successful strategies make cooperation contingent on behaviors of others in the group. To see how nondifferentiable fitness functions arise in such repeated games, consider the iterated n-person prisoner’s dilemma (or public goods game) [17–19], iterated *T* times in a life cycle. Social interactions of this kind are likely to be important in all kinds of social vertebrates, and especially primates. Chimpanzee patrolling and human food sharing may be examples. Suppose that individuals interact repeatedly in groups of size *n*, and the extent of individual prosocial action is a continuous variable (e.g. the amount of food shared, or the level of risk taken on). Let the value of this variable for individual *j* be *y_j_* and the fitness effect of one period of interaction for individual *j* be 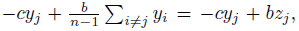 where the sum is over the other members of individual *j*’s group. Individual behavior is contingent. The wild types always give 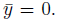 However there is a rare invading type that gives 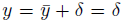 in the first interaction, where *δ >* 0 is small, and continues to give this amount as long as the other group members give in the average at least *δθ*, where *θ >* 0 is a threshold parameter. Setting 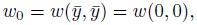 the fitness function takes then the following form:

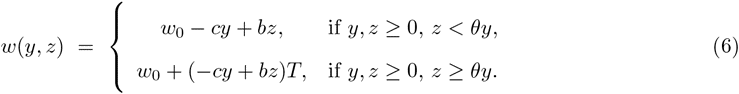

In order to compute the marginal fitness function *v*(*x*), using (3), we assume 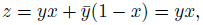, so that (6) becomes

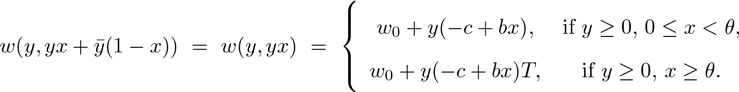

This yields, from (3), the non-linear marginal fitness function

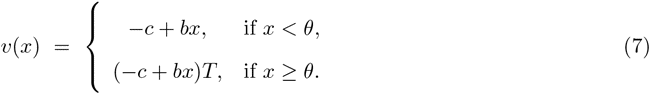

This expression for *v*(*x*) is intuitive, and can be obtained more directly from the following reasoning. If there is a fraction *x < θ* of mutants in the social environment, the focal actor will contribute *δ* and receive *δx < δθ* in the first iteration, and stop contributing. If, *x ≥ θ*, on the other hand, this actor will continue to contribute, as will all the other mutants in the group, for the same reason. In the former case the effect on the actors fitness is accounted as *−cδ* + *bδx*, while in the latter case it is this amount multiplied by the number of iterations, *T*. Since *v*(*x*) is the fitness effect divided by *δ*, we obtain (7) and its meaning becomes clear. The source of the non-linearity in *v*(*x*) in this example becomes also clear, as a consequence of the contingent behavior based on a threshold on the fraction of mutants in the group. When we compute *v*(*x*), this fraction *x* is fixed, and the resulting expression for *v*(*x*) will change in nature, as *x* crosses the threshold. Contingent behavior that leads to non-linear fitness functions is a common feature in the modeling of social evolution, especially of human cooperation; see, e.g., [20] and references therein.

It is important to understand that the non-differentiable *w*(*y, z*) of example (6) has well defined partial derivatives 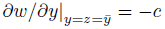 and 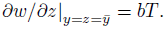 If one applied the Taylor-Frank method as proposed in [4–6], without knowing that differentiability is required, one would conclude that the condition for invasion of the mutant allele is given by (2), which here reads

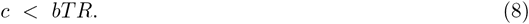

This would mean that if the game were repeated, say, 100 times in a life cycle, and *b* = 5*c*, then the critical relatedness required for the altruistic behavior to invade the population would be *R* = 0.002 = 0.2%; an extremely low value. We will see below that (8) is far from correct, and that a correct condition can be derived from (5), under biologically plausible assumptions.

Hamilton’s rule (2) is appealing because the only information needed about patterns of interaction is the relatedness *R*. Assuming *δ* small enough, *R* can be obtained from the distribution of neutral genetic markers in the same population. That is, there is a separation of time scales so that changes due to demographic processes occur much faster than changes due to selection. When (5) has to replace (2), *R* is not enough. More detailed information is needed about the distribution of *X*. This point was also made in [11]. However as long as selection is weak, the separation of time scales exists and the distribution of *X* can be calculated using the distribution of neutral genetic markers. Problems of this type have been addressed in a number of papers, including [15–17, 21]. This approach was applied in [17] to the iterated public goods game (6), in a population structured in groups, in which individuals compete within the groups, while groups also compete among themselves, and in which migration among groups also takes place. For this population structure, it was shown there that when groups are large the distribution of *X* is a beta(*α*,*β*) distribution, with parameters *α* = 1 and *β* = 2*×*(group size)*×*(migration rate) = (1*/R*)−1, which has probability density function *β*(1 − *x*)^*β*−1^, 0 *< *x* <* 1. The expected value in the generalization of Hamilton’s condition given by (5) can then be computed and this condition becomes

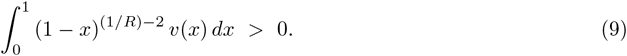

In the case of (6) and the corresponding (7), the integration in (9) is straightforward and the resulting invasion condition in terms of the threshold *θ* and the relatedness *R* was obtained and its consequences analyzed in the supplementary material of [17], Section 8. Here we recall only the case emphasized in that paper: when *θ* = *c/b* and *T* is large, the condition for invasion takes the simple form

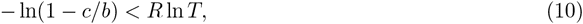

illustrating the usefulness of (5). This expression was used in [17] to show that even in large groups, strong altruistic behavior can proliferate when rare, under biologically realistic conditions, through population viscosity without the need of kin recognition, or greenbeard effects. For instance, when as above *T* = 100 and *b* = 5*c*, the critical relatedness required for the altruistic behavior to invade the population would be *R* = 0.048 = 4.8%. This is a modest value, compatible with available data for several species, [22] (Tables 6.4 and 6.5), [23], [24] (Table 4.9) and [20], but substantially different from the incorrect value obtained from (8).

It is important to understand that it is not the discontinuity in *v*(*x*) in the example above that is relevant, but rather its non-linearity. To illustrate this point, consider a generalization of (6) of the form

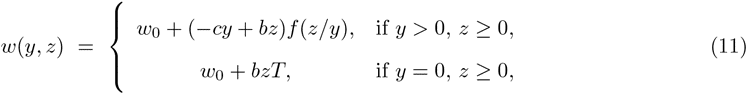

where lim*_x→∞_ f* (*x*) = *T*. Such a fitness function results, for instance, from a modification of the example above, in which the cooperative mutants cooperate in each iteration with a probability that may depend on the number of individuals that cooperated in previous rounds. For 0 *≤ × ≤* 1, the function *f* (*x*) gives the expected number of times that a cooperator cooperates, when the fraction of cooperators in its group is *x*. In case *f* is the threshold function that takes value 1 when *x < θ* and value *T* when *x ≥ θ*, (11) becomes (6). But one can argue that biological considerations, as the inclusion in the model of possible perceptual errors by the group members, suggest that an analytic function *f* would be more appropriate; possibly a sigmoid function that is close to 1 below the threshold *θ*, and close to *T* above this threshold. We will observe next that this makes no difference in our discussion. In this example, 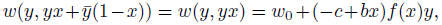 which yields, from (3), the marginal fitness function

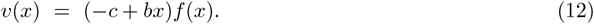

By expanding *f* (*x*) in powers of *x*, we see that *v*(*x*) is linear if and only if *f* (*x*) is a constant (otherwise *v*(*x*) would include powers of *x* larger than 1 in its expansion). This corresponds to the case in which the behavior of cooperators does not depend on the fraction of cooperators in the group. It is only in this very special case that (2) can be applied to example (11). But we can always apply (5) to this model, which for the population structure studied in [17] yields the invasion condition

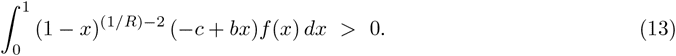

It is worth noting that if *f* (*x*) is a sigmoid function that is close to the threshold function that takes value 1 when *x < θ* and value *T* when *x ≥ θ*, then the invasion condition (13) is well approximated by the invasion condition for the model described by (6), which takes the form (10) when *θ* = *c/b* and *T* is large. In contrast, the application of the Taylor-Frank (2) to (11) produces also in this case the incorrect condition for invasion (8).

Another instance in which (5) was applied succesfully is [25], where we considered a population structure typical of human hunter-gatherers, with individuals organized in ethnolinguistic groups, divided into socially interacting bands. The distribution that we found for *X* in this case was more complicated then the beta distribution found in [17], but still simple enough to allow for estimates of critical values of relatedness needed for cooperation to proliferate. This allowed us to compare these theoretical threshold values of relatedness with available estimates of the levels of relatedness observed empirically ([20,23,26]) and in this way discriminate between mechanisms that could allow cooperation to spread from others that would not be viable. While certainly more complicated than (2), condition (5) can be analyzed in detail in some important cases, and provides transparent conditions for invasion of a rare social mutant into a monomorphic population, allowing for comparison between theory and data.

We conclude this section commenting on the paper [10], in which the Taylor-Frank methodology was applied in a situation in which the necessary differentiability is not satisfied. This paper considers groups of individuals that act in ways that depend on the phenotypes of all the individuals in the group. The fitness of each individual depends then on the actions of all the group members. Note that an instance of this is the iterated public goods game, with the strategies that we discussed above, corresponding to (6). From our discussion we know that differentiability fails in this case. Unaware of the limitations of the Taylor-Frank method, the authors of [10] used it (in the fashion that will be explained and criticized in the next section) in their equations (2) and (3). Their resulting borderline condition for evolutionary stability of a trait is given by their equation (5). In the case of the iterated public goods game discussed above, instead of the correct condition obtained from (5) (e.g., (10)), their condition reduces to the incorrect *c < bR*, independently of the number of iterations *T* and of the threshold *θ*. (For the computation of the various partial derivatives in the approach of [10], keep in mind that we are interested in the evolutionary stability of the equilibrium in which no one cooperates, i.e., all phenotypes are 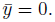 When *θ >* 0, in a large group, if only one individual is assumed to be a cooperative mutant, this individual will cooperate in the first round and not later, while the others will never cooperate. Using this observation for computing the partial derivatives at this point, we obtain then for the quantities defined in [10], *ρ* = 0, their *c* is identical to ours, while their *b* is ours divided by *N -* 1, where *N* is group size.) The fact that this paper was published in a leading journal in evolutionary biology, and that apparently their mistaken use of the Taylor-Frank method has not been noticed before highlights the relevance of pointing out the need of differentiability in this method, and of emphasizing the alternative solution when differentiability does not hold. (It is interesting that the direct application of the Taylor-Frank approach to (6) gives the invasion condition *c < bT R*, while its application in the fashion of [10] gives the invasion condition *c < bR*. Both are incorrect, but they are also different from each other. The reason for a difference between them is that (6) lumps together the effect of all cooperators in the group on the focal individual, while in [10] these are treated separately. Mathematically correct methods all produce the same answer to a given problem, but incorrect methods may produce various different answers to a single question, depending on the stage at which the mistake is made.)

## Regression coefficients cannot be replaced by partial derivatives

Suppose that a focal individual is chosen at random from the population. Define *y_•_*, *z_•_* and *w_•_* = *w*(*y_•_, z_•_*) as the random variables that are equal to the values that these quantities take for the focal individual. The invasion condition *w*^m^ − *w*^w^ *>* 0 can be rewritten in the following form, with definitions that are given below (see, e.g., [5, display 5], or [4, display 6.5]):

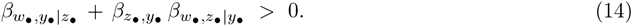

Here the three *β*’s are defined as the numbers that together with the proper choice of the constants *α^′^* and *α^″^* minimize

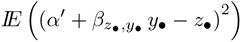

and

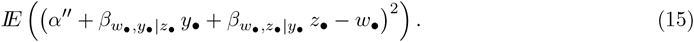

This definition in (15) says that 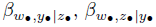 and *α^″^* are the numbers that make the function

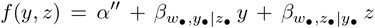

the best linear approximation to the function *w*(*y, z*), in the sense that it minimizes the square of errors weighted by probabilities over the values of *y* and *z*. (The definitions of the *β*’s can be rephrased in terms of projections of random variables on appropriate Hilbert spaces. For instance, the minimization of (15) is equivalent to the statement that the projection of the random variable *w_•_* onto the space of affine functions of *y_•_* and *z_•_* is of the form 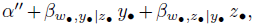 for some constant *α^″^*.) The condition (14) is appealing because 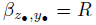 is the relatedness in the social environments, and therefore (14) is equivalent to

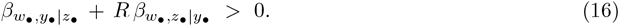

The *β*’s are usually referred to as regression coefficients, and we will use this terminology in what follows. As we explain next, if *w*(*y, z*) is differentiable, and the distribution of values of (*y_•_, z_•_*) is narrowly concentrated close to 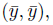 then the regression coefficients in (16) are the same as the marginal fitnesses derived using the Taylor-Frank method. Indeed, if *w*(*y, z*) is differentiable at 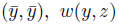 *w*(*y, z*) is well approximated by a linear function of *y* and *z*, in the neighborhood of this point. This means that

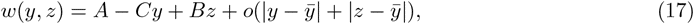

with 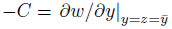 and 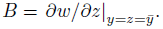 Thus

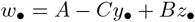

is a good approximation and hence the second optimization problem in (15) is solved by 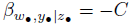 and 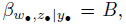 regardless of the details of the joint distribution of *y_•_* and *z_•_*. This approximation becomes better and better, as *δ →* 0, and therefore (14) is well approximated by Hamilton’s condition *C < BR*, in the limit of weak selection.

But suppose now that *w*(*y, z*) is not differentiable at 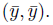 In this case, we can even add the assumption that the mutant types, with 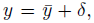 are rare, and still we will not have the approximate equalities between the regression coefficients 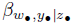 and 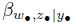 and, respectively, the partial derivatives 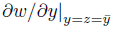 and 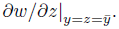 To illustrate this point, suppose that *w*(*y, z*) is given by (6). Recall that the partial derivatives at 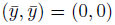 exist and are given by −*c* in the *y* direction, and *bT* in the z direction. Compare now the following two scenarios. (1) The distribution of (*y_•_, z_•_*) is concentrated in the region where *z < θy*. In this region *w*(*x, y*) is identical to the linear function *w*_0_ −*cy* + *bz*, and therefore, in this case, 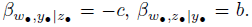 The distribution of (*y_•_, z_•_*) is concentrated in the region where *z ≥ θy*. In this region *w*(*x, y*) is identical to the linear function *w*_0_ + (−*cy* + *bz*)*T*, and therefore, in this case, 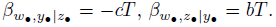 These two observations are true even when the distribution of (*y_•_, z_•_*) is concentrated mostly on 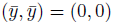 (rare mutants) and fully concentrated in a small neighborhood of this point (small *δ >* 0). It is then clear that the regression coefficients may be quite different from the partial derivatives even under these assumptions.

For another example, suppose that *w*(*y, z*) is given by Fig 1. The assumptions that we made about *δ* being small and the mutants being rare, implies that the distribution of (*y_•_, z_•_*) concentrates close to the point 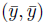 and on the segment 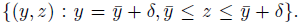 But the distribution over this segment depends on demographics—it is determined by the distribution of the random variable *X* that gives the number of mutants in the social environment of a mutant focal. Because *w*(*y, z*) is not well approximated by a linear function of *y* and *z* in the relevant region, (even for very small values of *δ*), the regression coefficients 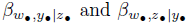 will depend on the distribution of *X* in a substantial way. To see why consider the function, *v*(*x*), shown in Fig 1. This function is very flat when *x* is close to 0 or 1, but is steeply increasing when *x* takes intermediate values. Now, compare three scenarios. (1) If the distribution of *X* is concentrated close to *x* = 0, then we will have 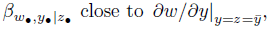 which is a negative number, and 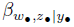 close to 0. (2) If the distribution of *X* is concentrated close to *x* = 1, then we will have 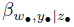 positive and again 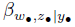 close to zero. (3) If the distribution of *X* is concentrated in intermediate values of *x*, then we will have 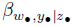 even more negative than 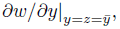 and 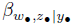 large and positive.

The idea that when selection is weak and mutants are rare (vanishing trait variation in the population) we would have in good approximation

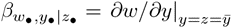

and

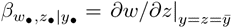

has been used to justify the Taylor-Frank method (e.g., [5, Box 6] and [4, as they justify deriving 6.7 from 6.5]). This idea is intuitive and appealing, but unfortunately it is not correct, unless *w*(*y, z*) is differentiable in the relevant region.

## Conclusions

Whether the Taylor-Frank method is appropriate to determine the stability of a monomorphic equilibrium against invasion by a rare social allele, with a small mutation, depends on the biological facts describing how the fitness of a focal mutant individual depends on the fraction *x* of individuals in its social environment that carry the same mutation. The method, as originally proposed in [6], can be properly applied only when this dependence is linear. This is not the case, e.g., in repeated *n*-person games. However, even when the Taylor-Frank direct fitness method is not appropriate as originally proposed, it can be replaced by the more general direct fitness expression (4), and the corresponding generalization of Hamilton condition for invasion (5).

## Acknowledgments

We are grateful to Clark Barrett for enlightening discussions. We are also grateful to Sam Bowles, Steve Frank, Anne Kandler, Laurent Lehmann and Jeremy Van Cleve for nice conversations and useful feedback on various aspects of this project and related subjects. This project was partially supported by the Center for Natural and Artificial Information Processing Systems of the University of São Paulo (CNAIPS-USP). No additional external funding has been received for this study.

